# Embedded 3D printing in self-healing annealable composites for precise patterning of functionally mature human neural constructs

**DOI:** 10.1101/2021.08.04.455135

**Authors:** Janko Kajtez, Milan Finn Wesseler, Marcella Birtele, Farinaz Riyahi Khorasgani, Daniella Rylander Ottosson, Arto Heiskanen, Tom Kamperman, Jeroen Leijten, Alberto Martínez-Serrano, Niels B. Larsen, Thomas E. Angelini, Malin Parmar, Johan U. Lind, Jenny Emnéus

## Abstract

Human *in vitro* models of neural tissue with controllable cellular identity, tunable microenvironment, and defined spatial arrangement are needed to facilitate studies of brain development and disease. Towards this end, embedded printing in jammed microgel supports (i.e., granular gels) holds great promise as it allows precise and programmable patterning of extremely soft and compliant tissue constructs. However, in contrast to the vast material landscape available for bulk hydrogels, granular printing support formulations are restricted to a handful of materials without the ability for facile adjustment of biofunctional properties of the cellular microenvironment. Therefore, there has been a need for novel materials that take advantage of versatile biomimicry of bulk hydrogels while providing high-fidelity support for embedded printing akin to granular gels. To address this need, we present a modular platform for bioengineering of neuronal networks via direct embedded 3D printing of human stem cells inside Self-Healing Annealable Particle-Extracellular matrix (SHAPE) composites. SHAPE composites consist of soft microgels immersed in viscous extracellular-matrix solution to enable precise freeform patterning of human stem cells and consequent generation and long-term maintenance of mature subtype-specific neurons that extend projections within the volume of the annealed support. The developed approach further allows multi-ink deposition, live spatial and temporal monitoring of oxygen levels, as well as creation of vascular channels. Due to its modularity, SHAPE biomanufacturing toolbox not only offers a solution for functional modeling of mechanically sensitive neural constructs, but also has potential to be applied to a wide range of biomaterials with different crosslinking mechanisms to model tissues and diseases where recapitulation of complex architectural features and topological cues is essential.

## Introduction

Astonishing architectural and operational intricacies of the human brain span multiple length scales to allow precise control of our cognitive and motor functions. Most of our understanding of this complex organ comes from a long tradition of studies using post-mortem tissue, animal models, and two-dimensional (2D) cell cultures. However, the pitfalls of these approaches have become increasingly recognized. For example, planar cell cultures do not recapitulate spatial complexity of the native tissue while results from animal experiments often fail to be translated to human trials due to interspecies differences^[1][2]^. Consequently, there has been a growing need for three-dimensional (3D) *in vitro* models that more faithfully recapitulate aspects of anatomical features and cellular processes underlying human brain development and disease. In addition to achieving authentic cellular identity, advancing neural 3D models requires controllable and defined spatial arrangement of cells within an extremely soft environment that mimics the native extracellular matrix (ECM) and promotes neuronal maturation, axonal outgrowth, and functional activity. In this regard, 3D printing holds great promise as a highly versatile biofabrication approach that has been leveraged to engineer *in vitro* constructs to mimic tissues such as heart, bone, muscle, and cartilage for disease modelling, drug screening, and regenerative medicine applications^[3][4][5][6]^. However, precise and programmable printing of neural constructs with defined 3D geometries remains particularly challenging due to the inherent lack of structural integrity exhibited by soft hydrogels.

A conventional solution to prevent deformations and collapse of mechanically weak engineered tissues is to print supporting strands from a stiffer secondary material that allows the final construct to retain the desired shape. Then, soft cell-laden hydrogel could either be printed between these strands^[7]^ or loaded into the construct post-printing^[8]^. However, this approach requires careful rheological finetuning of individual inks, has limited freedom of design inherent to traditional layer-by-layer deposition, and presents risk of material dehydration during longer printing times. As a powerful alternative, embedded 3D printing has emerged in the past decade^[9]^. Instead of depositing material on top of a surface in ambient air, in this paradigm-shifting approach, the ink is extruded through a syringe needle inside yield-stress support medium that is solid-like at rest but fluidizes in the vicinity of the moving needle tip to allow facile and precise ink deposition. Then, in the wake of the needle, the support resolidifies to keep the deposited material in place. While bulk hydrogels have been explored for embedded printing, they offer suboptimal rheological properties and restricted fabrication possibilities limited to a single layer^[10]^ or to vertical patterning^[11]^. For high-fidelity printing, solid-to-liquid transitions need to be smooth, spatially localized, rapid, and reversible^[12][13][14]^. Pioneering studies have demonstrated that jammed microgel assemblies (i.e. granular gels) fulfill these requirements and exhibit excellent rheological properties as supports for embedded printing of delicate structures from soft hydrogels and cells^[15][16]^. Since then, embedded printing has evolved into a versatile biofabrication platform used for the manufacturing of anatomically accurate tissue components^[17][18][19][20]^, patterning of cellular spheroids^[21][22]^, engineering of perfusable channels^[23][24][25]^, and investigation of cell-generated forces^[26]^.

Unfortunately, in contrast to the vast material landscape offered by bulk hydrogels, granular gels designed for embedded printing have been restricted to a handful of materials that allow simple chemical formulation and cost-effective production of microparticles in large quantities (e.g. gelatin, poly(acrylic acid), agarose). While efforts to tune the biochemical properties of the constituent microparticles to mimic tissue-specific ECM could broaden embedded bath functionality, the related expense and technical challenges practically eliminate these advantages. Furthermore, traditional granular supports immersed in low viscosity liquids such as cell culture media, water, or saline require tight packing of microparticles to allow high-fidelity printing. With maximized microgel volume fraction, the presence of the continuous material in the interstitial space (i.e., the void surrounding the particles) is minimized and with it the ability of adding biofunctional moieties to the support. Instead, ECM-based polymers are mixed into the ink with cells to provide tissue-specificity. However, this approach confines cell-ECM interactions exclusively to the extruded volume thus impeding neuronal tissue growth and formation of extended neuronal projections between features of the engineered constructs. Consequently, there has been a need for novel printing supports that leverage the diversity of bulk hydrogel formulations while providing rheological benefits of granular gels. Towards this end, methods for formulating composites of the two types of material must be developed to allow straightforward control over cellular microenvironment together with precise patterning of cells in 3D. While composite materials have been explored for tissue repair applications they remain elusive as embedded 3D printing supports^[27][28][29]^.

Here, we present a modular approach for simple generation of hybrid materials termed **S**elf-**H**ealing **A**nnealable **P**article-**E**CM (SHAPE) composites. SHAPE supports consist of hydrogel microparticles with increased interstitial space immersed in viscous biofunctional polymer solution. As such, they provide both the physical support for high fidelity embedded printing and the cell-interactive microenvironment for healthy cellular growth, maturation, and activity. In contrast to traditional granular gels, the expanded presence of the continuous phase in SHAPE composites allows exploration of the broad spectrum of materials already established for bulk hydrogel systems. In particular, we formulated SHAPE support from thermally crosslinkable cell-adhesive ECM polymers and cell-inert alginate microparticles to facilitate precise embedded printing of human neural stem cells (hNSCs). Then, we demonstrated that annealing of the support due to crosslinking of ECM polymers creates a favorable 3D microenvironment for the generation, long-term maintenance, and maturation of subtype-specific human neurons. Importantly, annealed SHAPE support promotes omnidirectional axonal pathfinding and elongation into previously uninhabited support volume leading to the development of neuronal constructs with delicate neuronal morphology and mature functional activity. The presented approach furthermore provides a solution for the engineering of vascular channels and temporal 3D mapping of oxygen concentrations thus giving rise to a versatile biomanufacturing toolbox that can employ a wide range of biomaterials and crosslinking mechanisms with potential beyond neural tissue modelling applications.

## Results

SHAPE support material consists of soft hydrogel microparticles (the granular component, ~70% volume fraction) entangled in a viscous polymer solution (the continuous component, ~30% volume fraction) (Fig. 1A). To mimic the native microenvironment, we formulated the polymer solution from purified and recombinant ECM macromolecules (collagen, laminin, hyaluronic acid, and fibronectin). While at 4°C and physiological pH it behaves as a viscous liquid, this polymer mixture undergoes gentle gelation at 37°C due to thermal crosslinking of collagen. As such, the ECM-based continuous component is designed to promote differentiation of hNSCs of ventral mesencephalic origin into midbrain neurons containing dopaminergic subpopulation (their natural developmental trajectory) (Fig. S1). To create the granular component, we adapted a protocol for the generation of microparticles via mechanical fragmentation of internally crosslinked alginate hydrogels^[14]^. In comparison to other tested methods that yield microparticles of appropriate size (Fig. S2), this approach allows production of large quantities of alginate microgels without the need for specialized equipment or harmful reagents. By utilizing internal gelation mechanism, hydrogel is crosslinked uniformly resulting in high transparency of the final product. Furthermore, alginate granular gel supports have been successfully used for embedded printing making them an excellent reference for the rheological characterization of SHAPE material formulation ^[18]^.

**Fig. 1.**
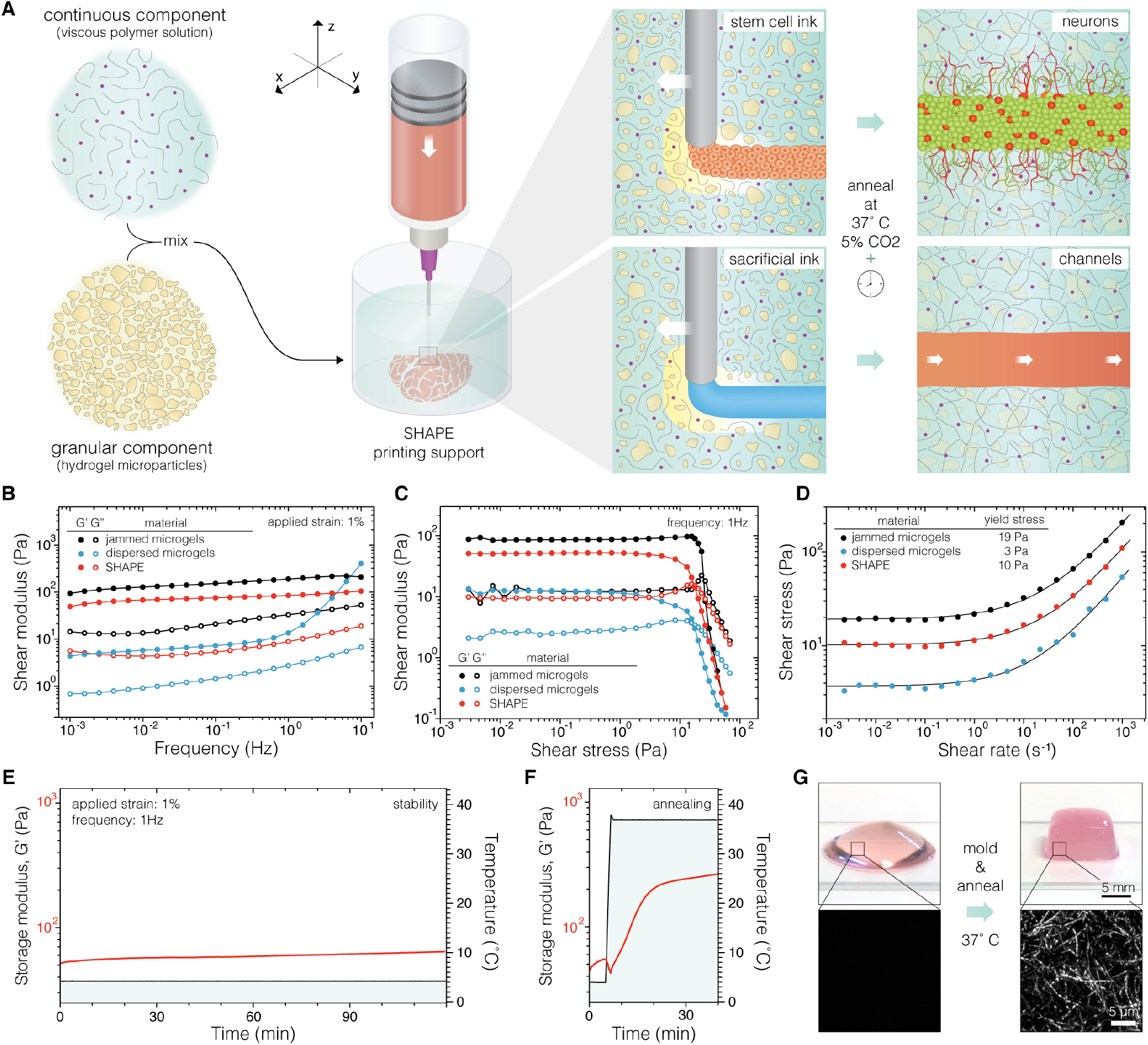
The methodological concept and rheological characterization. (**A**) Graphical illustration of the printing process in SHAPE support bath. (**B**) Storage (G’) and loss moduli (G”) of jammed microgels and SHAPE material show weak frequency dependence while dispersed microgel system exhibits drastically lower moduli with strong frequency dependence. (**C**) Measurements of shear moduli as a function of applied strain amplitude show solid-to-liquid transition. (**D**) Yield stress of the formulated material systems is determined by fitting a classic Herschel-Bulkley model to shear stress values measured at many constant strain rates. (**E**) Stability of storage modulus for SHAPE material support at 4°C over a period of two hours. (**F**) Increase in storage modulus when temperature is raised from 4°C to 37°C indicates temperature-induced collagen crosslinking. (**G**) The top row displays photographs of SHAPE hydrogel before (left) and after (right) annealing and molding. Corresponding reflectance microscopy images in the bottom row show the formation of collagen bundles in the annealed support.

The ability of granular gels to behave as solids at rest and to reversibly liquify at low shear-stresses relies on jamming transitions of microparticles at high packing density. This liquid-like solid characteristic is lost at lower packing densities as particles become separated. In turn, with reduced interparticle contact and interaction, the physical properties of the system become increasingly dependent on the composition of the continuous phase. We investigated this effect by replacing the polymer solution with cell culture media (Fig. 1, B-D). In comparison to reference values set by tightly jam-packed microgels (~100% particle volume fraction), dispersing alginate microparticles in Newtonian liquid with low viscosity such as cell culture media leads to impaired rheological behavior with 30-times lower shear moduli and frequency-dependent instabilities. However, when the interstitial space is filled with viscous ECM polymer solution (non-Newtonian pseudoplastic material ^[30]^) to form SHAPE composite support, favorable rheological properties of the system are restored. Like tightly packed granular gels, SHAPE composite responds to low stresses as an elastic solid while exhibiting liquid-like behavior when yielded. We determined the yield stress of the hybrid material at 4°C to be around 10 Pa, a value within optimal range for embedded printing of cellular inks ^[31]^. At this temperature, the SHAPE composite retains rheological characteristic for at least two hours (shear storage modulus increased by 14 Pa), indicating its suitability for longer printing times (Fig. 1E). Then, when temperature is increased to 37°C, thermally induced self-assembly of collagen into physically crosslinked fibers leads to annealing of the material allowing it to hold a predefined shape (Fig. 1, F and G).

To test how the formulated ECM-based composite performs as a support material for embedded 3D printing of cellular inks, we loaded hNSCs from the human ventral midbrain stem cell line ^[32]^ in a syringe attached to a 3D printer and extruded them along a predefined path in three different support materials (Fig. 2A). As expected, printed structures in the jam-packed microgel support were reproduced with high fidelity. On the other hand, dispersed microgels in cell culture media presented a poor environment for embedded printing. Due to the lack of spatial localization of the solid-to-liquid transition, printed layers are disturbed when their location is revisited by the needle in consecutive printing planes (wavering lines in the woodpile design). Furthermore, when the needle comes in proximity of the previously printed structure too soon, due to slow resolidification of the support, liquified regions expand causing the printed features to be dragged in the direction of the printing path. This effect is particularly intensified in designs with a dense infill (warping of the square construct). When printing inside SHAPE hydrogel, the stability of the process is restored indicating that the introduction of ECM polymers to the expanded interstitial space results in spatially localized thixotropic responses at sufficiently small timescales to allow for precise ink deposition and rapid self-healing of the support in the wake of the needle.

**Fig. 2.**
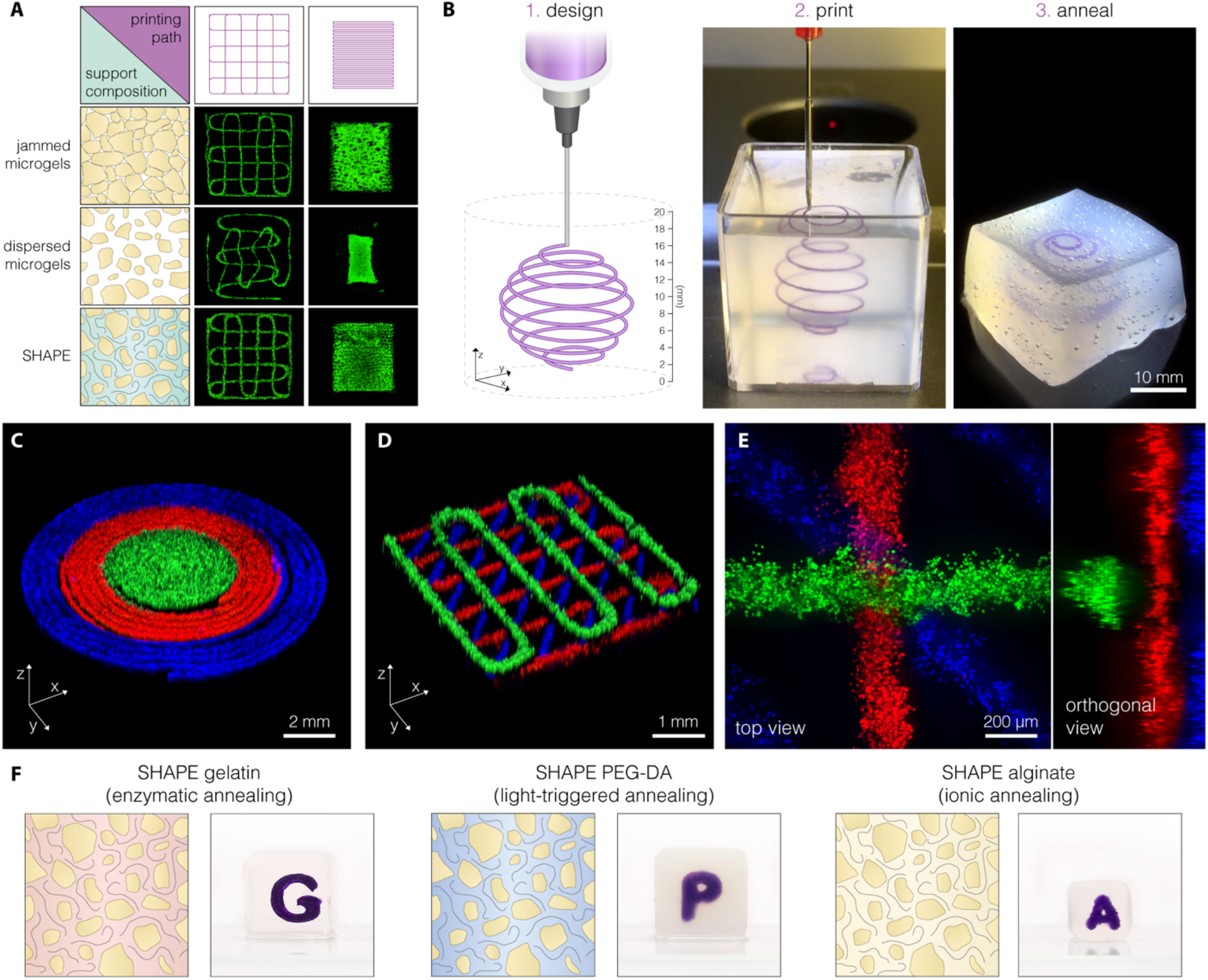
Embedded 3D printing in SHAPE hydrogel support. (**A**) Fluorescent images of hNSCs labelled with Calcein-AM show the fidelity of embedded printing of a programmed path (purple, top) supports with varying packing density and continuous phase composition (left, blue). (**B**) Example of a printing process showing omnidirectional printing capability. Polystyrene beads were used in the ink instead of cells to visualize the structure. (**C**) Concentric rings printed from three hNSC inks. (**D**) Stacked crosshatch structure printed from three hNSC inks. (**E**) Magnified view of a junction in a stacked crosshatch structure showing intact layers clear vertical separation. Distinct hNSC inks were created by staining live cells with Calcein-AM (green), Calcein-Red-AM (red), and Hoechst (blue). (**F**) Alternative SHAPE formulations provide diverse annealing mechanisms. While keeping the same granular component, we varied the contents of the continuous phase: 10% gelatin (left), 30% PEG-DA (middle), and 1% alginate (right). The images show annealed SHAPE supports displaying structural stability with visible printed constructs inside of each hydrogel.

To further explore the possibilities of embedded printing in the developed composite material, we generated several structures that showcase different aspects of the approach. Firstly, we created a spherical spiral with a 15 mm diameter in a continuous freeform manner (Fig. 2B, movie S1). Once printing was finished, the entire volume (16.5 cm^3^) of the composite support was successfully annealed to form a hydrogel block that can be removed from the container. The printed spiral remains embedded inside the free-standing hydrogel displaying structural integrity achieved by annealing. Then, to investigate the ability to create tissue constructs from multiple cellular inks using the developed approach, we loaded three syringes with living hNSCs (from the same cell line) containing different fluorescent markers. Cellular inks were printed in consecutive manner (green, red, and then blue ink). The results showed that embedded 3D printing in SHAPE hydrogel support was suitable for the precise generation of both planar structures with defined regionalization (Fig 2C) and stacked structures with delicate features (Fig. 2D). The fabricated constructs displayed high fidelity without visible offsets or damages caused by the reentry of the needle during ink exchange. Upon closer examination of the stacked layers (~20 μm separation), we did not observe mixing of the cells from adjacent layers indicating that needle movement within 100 μm of the printed structure did not cause material perturbations that were able to notably distort the preexisting features (Fig. 2E).

It is important to note that the underlying principles that make SHAPE material an excellent support system for embedded 3D printing are not restricted to the formulation presented in this work and can be easily adapted to other tissue types by tuning the contents of both the granular and continuous component. While here we formulated the continuous component to allow thermal annealing of the support and promote hNSC differentiation in 3D, its contents can be exchanged to enable different crosslinking mechanisms or to introduce novel biofunctional moieties. As an example, we replaced ECM polymers (while keeping alginate microgels in the granular component) with gelatin, poly(ethylene glycol)-diacrylate (PEG-DA), or alginate to respectively allow enzymatic, light-induced, or ionic crosslinking of the system while still providing support for high-fidelity embedded printing (Fig. 2F, S3). The modularity and flexibility of SHAPE hydrogel system thus opens new possibilities for material exploration in embedded 3D printing.

After printing hNSCs in the ECM-based composite, the support was immediately annealed and then the cells were differentiated for two months. During this time, the programmed geometry (crosshatch design with 1 mm spacing between 200 μm thick lines) was preserved with defined features both within and between printing planes (Fig. 3, A, B, and E). Microscopic analysis of the patterned lines revealed dense 3D cellular architecture, a result of hNSCs self-organization, and thin projections emanating from cell bodies that effectively penetrate the bulk of the hydrogel in all directions (Fig. 3, C and D, movie S2). Migratory cells were not observed in the spaces between lines. These findings suggest that the microenvironment provided by annealed SHAPE composite is permissive for axonal pathfinding and growth. Similar findings were observed in a square construct with a dense infill (Fig. S4). Clear separation of the differentiated construct into the somatic compartment (extruded volume with cell bodies) and the axonal compartment (initially empty hydrogel volume progressively populated by neural projections) is one of the prerequisites for the generation of *in vitro* models with architectural and functional relevance. In fact, the anatomy of the brain is intertwined with clusters of neurons that, besides communicating with each other to form local microcircuitries, extend their projections outward to form interregional axon tracts. The developed printing approach presented here would not only allow regulation of local cell densities (through patterning design and cellular concentration of the ink), but also offer a possibility for the regulation of neurite outgrowth and axonal pathfinding.

**Fig. 3.**
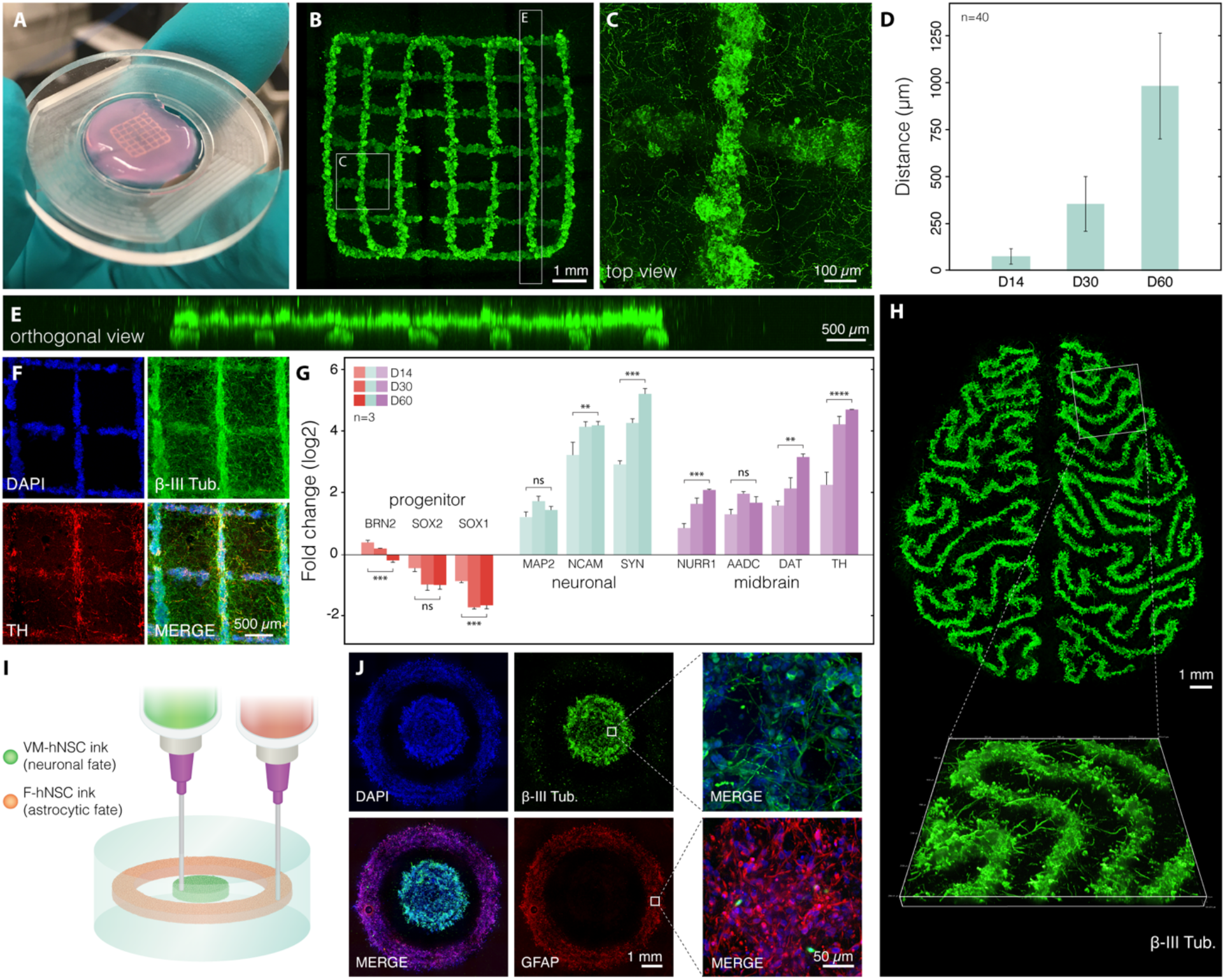
Generation of authentic human neurons within 3D printed constructs. (**A**) Photograph of a 3D printed construct in annealed SHAPE support after 2 months of differentiation. (**B**) Confocal image of the same construct labelled with Calcein-AM. (**C**) Maximum intensity projection of a 200 μm optical section showing that neural projections extend in the volume between printed lines. (**D**) Quantification of the extent of projection outgrowth away from a 3D printed structure as a function of the differentiation time. (**E**) Orthogonal projection showing that the structural integrity with defined features in different layers are preserved. (**F**) Fluorescence images of immunostained 3D printed construct after 2 months of differentiation displaying neuronal and dopaminergic markers (β-III tubulin and TH respectively). (**G**) RT-qPCR gene expression analysis of the printed construct at 3 different time points indicating midbrain patterning and neuronal maturation. Non-differentiated hNSCs were used as a reference. (**H**) Maximum projection fluorescence image of neurons in a 3D printed construct resembling brain meanders with a 3D close-up view of a selected segment. (**I**) Illustration showing the printing of cells with orthogonal differentiation trajectories by using 2 inks containing hNSCs with different predefined fates. Inks were printed in consecutive order in concentric circle design within the same SHAPE support. The outer radius of the inner and outer disk is 2 and 4 mm respectively. (**J**) Fluorescence images of fixed and immunolabelled constructs after 1 month of differentiation. DNA counterstain shows the presence of cells both in the inner and outer ring but staining for GFAP and β-III tubulin reveal distinct protein expression characteristics between cells generated from each ink. Cells display elongated morphology indicating successful differentiation.

In addition to exhibiting neuronal morphological phenotype, the differentiated cells expressed characteristic neuronal and midbrain markers in line with their fetal ventral mesencephalic origin. In contrast to non-annealed granular gels, structurally stable SHAPE hydrogels protect the delicate features during the process of fixation and immunolabelling, therefore allowing not only morphological investigation of live cells but also detection and visualization of antigens in fixed samples. Immunocytochemistry revealed that printed constructs are rich with cells expressing β-tubulin III (cytoskeletal protein found almost exclusively in neurons) with a subpopulation expressing tyrosine hydroxylase (TH, rate-limiting enzyme in the production of dopamine) (Fig. 3F). Furthermore, real-time quantitative PCR (RT-qPCR) analysis demonstrated time-dependent decrease in expression levels of neural progenitor markers (*BRN2*, *SOX1*, *SOX2*) with simultaneous increase in the expression level of neuronal markers (*MAP2*, *NCAM*, *SYN*) and markers associated with dopaminergic neurons (*NURR1*, *AADC*, *DAT*, *TH*), confirming progressive maturation of dopaminergic neurons with midbrain identity (Fig. 3G). Importantly, we demonstrated that successful hNSC differentiation is not restricted to simple geometries but can be expanded to 3D printed structures with arbitrary designs, such as meandering folds resembling brain gyrification emphasizing the flexibility of embedded printing as a powerful biofabrication technique (Fig. 3H).

One of important aspects in disease modeling and tissue engineering is the ability to incorporate multiple cell types within an *in vitro* construct. To achieve this, stem cell populations with orthogonal developmental trajectories could be printed as a potential way to model interactions between different cell types in the brain. To showcase the proposed principle, we created two distinct stem cell inks. In one syringe, we loaded hNSCs described above (with ventral mesencephalic origin). In the second syringe, we loaded hNSCs dissected from a forebrain region that have the propensity to differentiate into astrocytes ^[33]^. The two stem cell populations were printed consecutively as concentric rings, which were differentiated for 4 weeks (Fig. 3I). Immunocytochemistry of the differentiated construct showed a clear distinction between the cell types generated from the two cellular inks within the same support. While the inner disk consists mainly of neurons (marked by β-tubulin III), cells in the outer ring show high expression of GFAP, intermediate filament protein present in astrocytes (Fig. 3J). Embedded printing displayed here of distinct stem cell inks could allow researchers to investigate the importance of spatial arrangement in the interaction between cell types.

To further validate the ability of the annealed ECM-based SHAPE support to provide favorable environment for neuronal maturation and functional activity, we performed calcium imaging and electrophysiological measurements. Global presence of intracellular calcium waves in the absence of external membrane-depolarizing stimuli indicated spontaneous neuronal signaling within 3D networks, which is a sign of a healthy neuronal population (Fig. 4A). Then, by using a microelectrode for single-cell electrophysiological recordings, we targeted cells labelled with synapsin-GFP that display neuronal morphology (Fig. 4B). Cells analyzed after two months in culture by whole-cell patch-clamp technique showed capability to fire current-induced action potentials (APs) (Fig. 4C) and presented inward sodium and delayed rectifying potassium currents (Fig. 4D), indicating presence of voltage-gated sodium and potassium channels on their membrane and a neuronal functional profile. Most cells expressed a hyperpolarized resting membrane potential (Fig. 4E) similar to endogenous dopamine neurons. While a proportion of analyzed cells (n = 6) exhibited mature AP, other cells demonstrated immature AP (n = 5) or no firing of AP (n = 11) suggesting different stages of differentiation into functional neurons (Fig. 4F). Together, the data from functional characterization provided evidence that hNSCs within the 3D printed constructs can differentiate into healthy and functionally mature neurons.

**Fig. 4.**
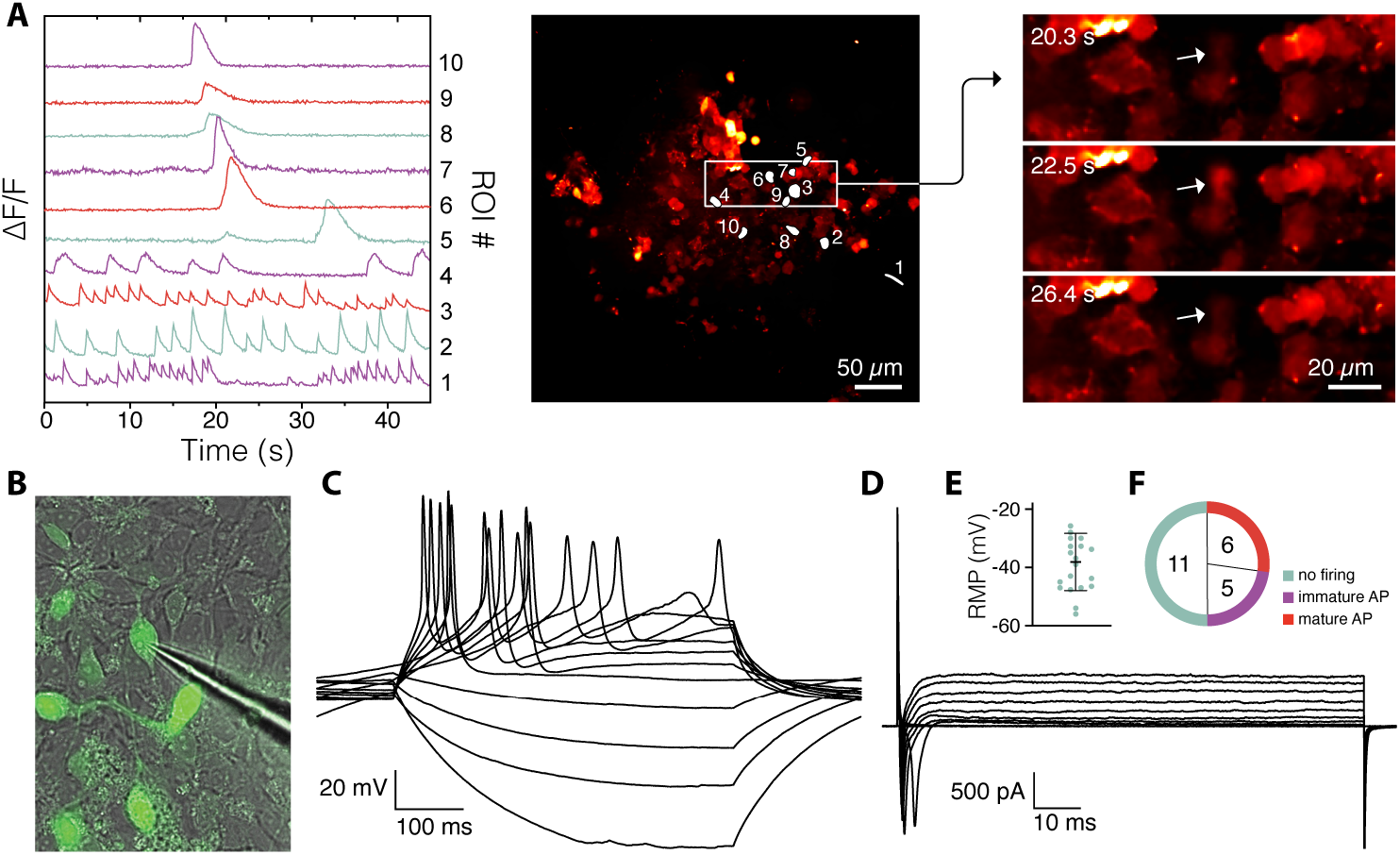
Functional assessment of neuronal activity. (**A**) Differential fluorescence intensity profiles showing changes in intracellular Ca^2+^ as a function of time (left); regions of interest marked for intensity profiling (middle); selected timeframes displaying Ca^2+^ transient in a cell marked by an arrow (right). (**B**) Image of a microelectrode targeting the synapsin-GFP neuron during whole-cell patch-clamp recordings. (**C**) Representative trace of whole-cell patch-clamp recordings performed in current-clamp mode displaying induced action potentials (APs) elicited by a neuronal cell (**D**) Representative trace of whole-cell patch-clamp recordings performed in voltage-clamp mode showing the presence of inward sodium/outward potassium currents in the targeted cell. (**E**) Resting membrane potential (RMP) measured during electrophysiological recordings demonstrating a hyperpolarized state, typical of neuronal cells. (**F**) Analysis of APs elicited by recorded cells (n = 22).

A major challenge in 3D tissue modeling and tissue engineering is the control over tissue construct oxygenation. Optimal oxygen supply to the cells is important not only for their survival (e.g., poor oxygenation of thick tissue constructs leads to the formation of a necrotic core) but can also interfere with the differentiation process, leading to undifferentiated cells or erroneously differentiated into unwanted cell types. We integrated a non-invasive optical method for oxygen sensing based on quenching of phosphorescence using microbead sensors embedded in and around the 3D printed cellular construct. This recently developed method allows 4D mapping of oxygen tension in cellular microenvironments using confocal phosphorescence lifetime microscopy ^[34]^. Oxygen sensitive microbeads can either be directly extrusion-printed in the support or mixed with the cellular ink during printing (Fig. 5A). SHAPE hydrogel support keeps microbeads in place after embedding thus allowing for controlled 3D distribution of oxygen microsensors. Oxygen maps were acquired between day 2 and day 50 to map both the spatial 3D distribution of oxygen tension in the neuronal constructs and the progression of oxygen tensions and gradients over time. A simple 3D printed disk exhibited oxygen tensions between approximately 2% dissolved O_2_ (DO) up to a radius of 2 mm from the center and increased to approximately 6% on the edges of the printed tissue construct (Fig. 5B). Oxygen levels gradually increase in the volume surrounding the disk confirming that low oxygen tension within the printed construct is caused by cellular consumption. Next, we explored the ability to tune oxygen levels through blueprint design. We printed a rectangular construct with varying infill density (increasing line spacing from left to right) (Fig. 5C). The cellular construct exhibits oxygen gradients along the direction of ascending line spacing from approximately 5.6%-16.1% DO (Fig. 5D). The investigation of the influence of infill density and extrusion rates on oxygen tensions within the printed constructs indicate linear relations for the observed parameter ranges further reinforcing the idea of design-driven control over tissue oxygenation (Fig. 5, E and F). The monitoring of oxygen tension over time in constructs with high infill density revealed an approximately linear decrease in DO, from 8.6% to 1.7%, during the first two weeks of culture when undifferentiated stem cell can still divide, after which the oxygen tensions remained stable at 1.7% until the end of the monitored 30-day period, pointing to the establishment of a postmitotic cell population (Fig. 5G). These results show that oxygen concentrations and gradients can not only be measured spatially in 3D but also at arbitrary intervals over time, allowing another level of control over tissue construct development. This could for example be used to time a switch from growth to differentiation media when desired oxygen microenvironments are achieved. This slow establishment of desired microenvironments can ease the cellular stress from fast and strong changes in oxygen concentration after seeding thus allowing differentiation in a stable microenvironment.

**Fig. 5.**
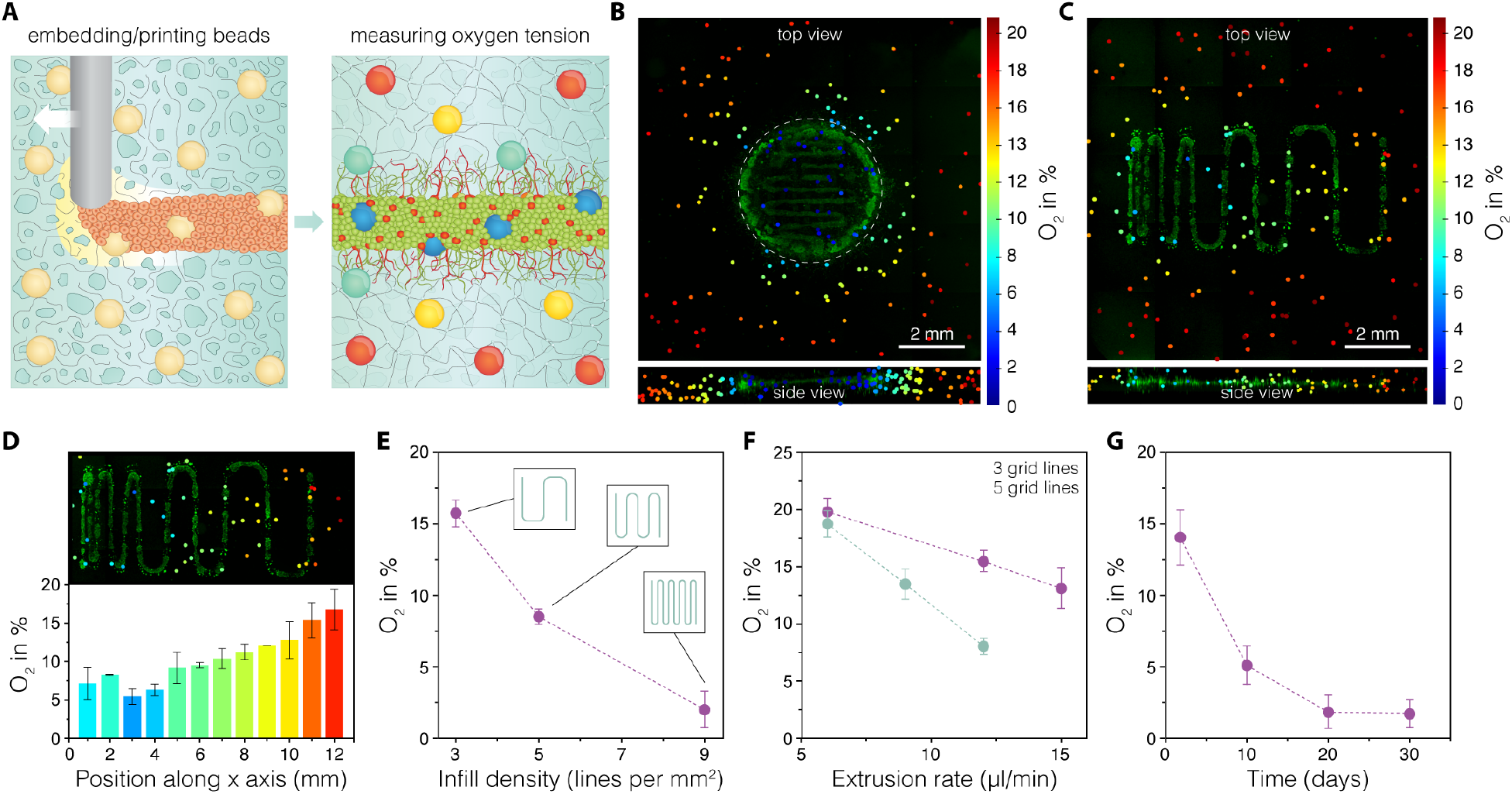
Oxygen mapping in 3D printed neuronal constructs. (**A**) Graphical illustration depicting the 3D oxygen mapping approach. Oxygen sensitive polystyrene microprobes are integrated in the printed cellular constructs both by mixing with the SHAPE printing support and by mixing with the stem cell ink. Oxygen levels are measured through optical readout based on oxygen-mediated quenching of phosphorescence. (**B**) Simultaneously acquired 3D oxygen and live-cell image stacks of a 3D printed cellular disk with collapsed Z dimension (top) and collapsed projection of 18 vertical planes extracted through a vertical center axis in rotational steps of 20 degrees (bottom). (**C**) Oxygen map of 3D printed neuronal construct with ascending grid spacing with collapsed Z-dimension (top) and collapsed side view of the tissue construct area (bottom). (**D**) Patterning induced oxygen gradient in the construct with ascending grid spacing. (**E**) Influence of infill density on the oxygenation of the 3D printed construct. (**F**) The effect of line thickness (extrusion rate) on the construct oxygenation at two different infill densities. (**G**) Monitoring of oxygen tension over time.

Embedded 3D printing of sacrificial structures presents a viable approach for the generation of vascular channels, a crucial aspect of every tissue engineering toolbox. Here, we directly extruded gelatin ink maintained at 50°C through a 500 μm blunt needle into cold SHAPE hydrogel support bath to create a 65 mm long meandering structure (Fig. 6A). Upon deposition, the ink rapidly solidified allowing for effective printing of arbitrary designs. In contrast to the composite support that stiffens during the annealing process (50°C, 5% CO_2_), gelatin structures melt. Once liquid, gelatin can be displaced from the support material by injecting warm cell culture media, leaving open channels in place of the sacrificial structure (Fig. 6B). The printing process generates channels with round cross section (Fig. 6C) and uniform size along the length of the channel (Fig. 6D). Finally, we printed a channel design with branching points to showcase the ability of the sacrificial ink to fuse at points of contact during printing (Fig. 6E). Infusion of colored liquid into the channels showed simultaneous progression of the dye through parallel channels after branching, indicating symmetry in the branching point shape and generated back pressure, verifying that branched channels were manufactured without constrictions or size differences at contact points during printing (movie S3).

**Fig. 6.**
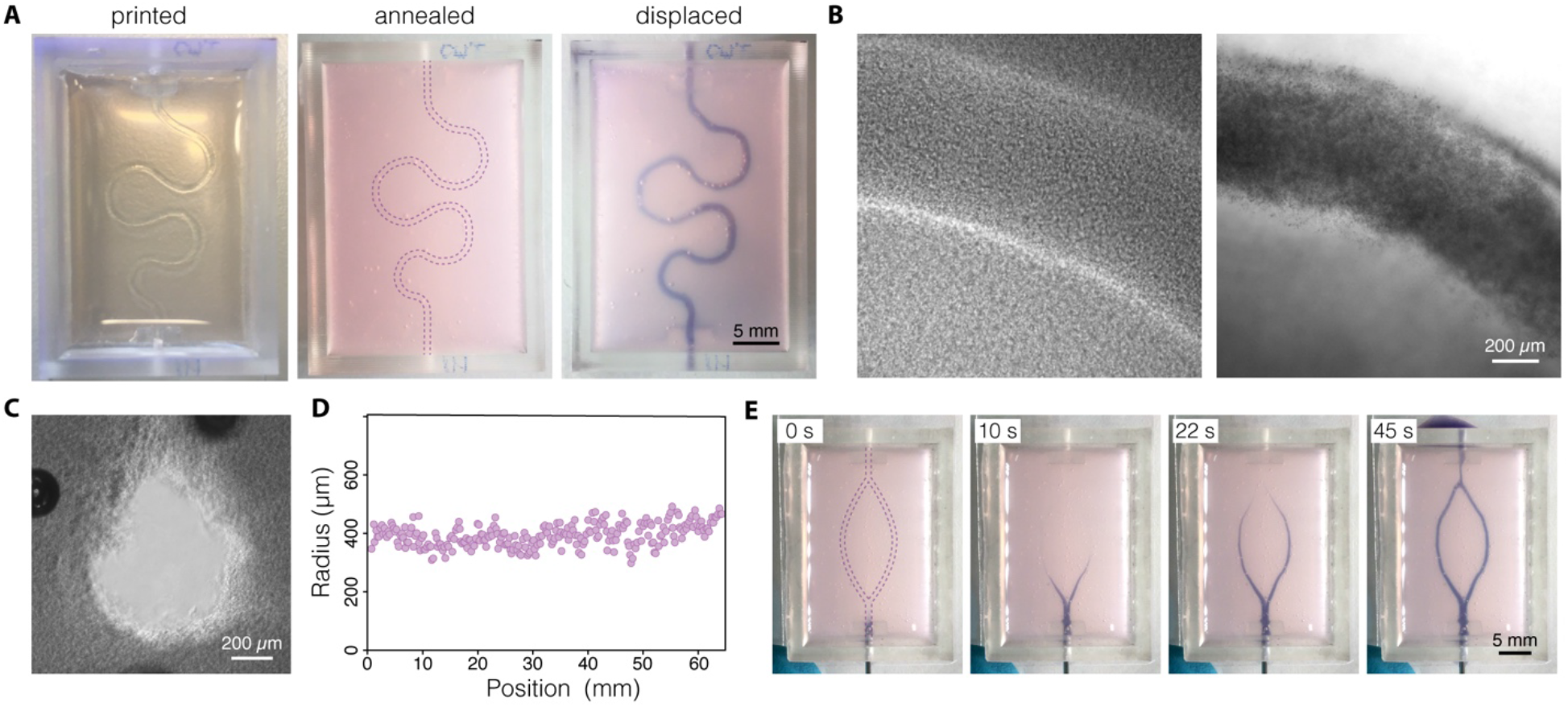
Printing vascular channels inside SHAPE material support. (**A**) Images showing the process of generating channels by printing and evacuating sacrificial ink: solid sacrificial gelatin structure visible immediately after printing (left), annealed support (middle), liquid sacrificial gelatin displaced and filled with a dye (right). (**B**) Channels before (left) and after (right) displacement of the sacrificial gelatin structure as seen under brightfield microscope. Channels are filled with polystyrene beads for visualization. (**C**) Channel cross section under phase contrast microscope displaying collagen fibers in the bulk around the channel. (**D**) Direct writing of sacrificial gelatin ink produces printed strands with uniform dimensions as indicated by the measurements of channel radius along the printed path. (**E**) Time-lapse images exhibiting perfusion of a branching sacrificial structure show simultaneous gelatin displacement of both channel branches indicating successful strand fusion at the branching point.

## Discussion

SHAPE material system provides a unique methodology for the formulation of supports for embedded 3D printing of cellular inks, soft materials, and channels that addresses the need for a versatile bioengineering route for the modeling of brain tissue. The modular nature of the technology (interchangeable granular and continuous components) enables exploration of a wide biomaterial landscape that has up to now been mostly available for conventional bulk hydrogel cultures. In the hydrogel precursor state, the rheological properties of the composite support can be tuned to allow freeform 3D biomanufacturing of multi-ink constructs with delicate features and truly arbitrary design. In parallel, biofunctional and mechanical properties of the annealed composite hydrogel can be tuned to mimic native cellular microenvironments that provide a 3D substrate for stem cell differentiation into a desired cell type (e.g., dopaminergic neurons presented in this work). We showed that subtype-specific neurons were successfully generated in the 3D printed cellular constructs and maintained for at least 2 months. Neurons acquired native-like morphology, mature functional activity and gene expression profile. Furthermore, they seamlessly integrated in the ECM network, that formed around the printed structures during the annealing of the hydrogel support, by extending axonal projections away from the somatic regions of the printed constructs and into the bulk of the composite support. These findings support previous claims that composite materials are able to promote connectivity of bioengineered neuronal networks ^[35]^. SHAPE biomanufacturing platform could also provide new opportunities towards neural circuit modeling where topological cues, the precise spatial organization and communication of defined cell types, are essential to the recreation of the native tissue architecture. For example, classification of midbrain dopaminergic neurons into A9 and A10 populations can currently only be determined through their anatomical location and region-specific innervation potential (dorsolateral striatum for A9 and nucleus accumbens and prefrontal cortex for A10) in animal models of Parkinson’s disease ^[36]^. Together with the ability for long term culture, embedded printing could be used to recreate native topological cues *in vitro* to study neural circuit formation and slow developing human pathological phenotypes. Brain organoid technology, a 3D *in vitro* culture approach that relies on the self-organizing ability of differentiating pluripotent stem cells to recapitulate neurodevelopmental processes in a physiologically relevant environment, could be combined with the embedded printing approach to create a powerful toolset for the next generation of spatially defined and automated brain tissue models ^[37][38][39]^. In our work, 3D oxygen mapping allowed manipulation of oxygen tension through construct blueprint design leading to the establishment of important printing parameters of infill density and extrusion rate for desired tissue oxygenation level. Thus, oxygen microenvironments that mimic *in vivo* brain oxygenation (1-3.5% DO) can be created ^[40][41]^. This factor is seldomly validated in 3D tissue models even though physiological oxygen level has a strong effect on stem cell differentiation and neuronal metabolism *in vitro* ^[42][43]^. In addition, printing parameters to achieve not only single oxygen concentrations but oxygen gradients in tissue constructs mimicking *in vivo* gradients could be validated and established. This method thus allows automated, high throughput establishment of organ models with 3D oxygen gradients, likely also exhibiting cellular signaling gradients. Finally, channels could be engineered within the 3D printed tissue models to provide nutrients and oxygen inside tissue constructs with dimensions and cell-density that exceeds capabilities of design-based oxygen tension manipulation ^[24]^. Furthermore, vascular channels could be leveraged to actively maintain gradients of patterning factors and chemo attractants/repellents to respectively form regionalized brain organoid models and actively guide axonal pathfinding ^[44][45]^. In conclusion, embedded 3D printing in SHAPE supports offers a versatile and flexible approach for functional modeling of mechanically sensitive tissues such as brain tissue. Printing support formulation presented here leverages affordable materials and simple manufacturing techniques but at the same time provides basic principles that will allow other research groups to utilize a range of natural or synthetic materials, crosslinking methods, and microparticle generating approaches in future applications of embedded 3D printing.

## Materials and Methods

### Preparation of the composite support material

1% sodium alginate (80-120 cP, Wako) was dissolved in sterile water for 4 hours at 60°C with constant stirring and filter sterilized (0.45 μm pore size). Then, equal volume of sterile alginate solution and 2 mg ml.^−1^ CaCO3 in sterile water were mixed and stirred at room temperature for an hour on a magnetic stirrer. This leads to 0.5% alginate and 1 mg mL^−1^ CaCO_3_ solution to which acetic acid was added in 1:500 ratio and stirred overnight at 650 RPM to allow crosslinking of alginate. Alginate particles generated this way were then homogenized at 15000 rpm for 10 min with T25 digital ULTRA-TURRAX homogenizer (IKA) equipped with S25N-10G dispersing tool to mechanically break down hydrogel into microparticles. Microgels were centrifuged at 18500 G for 20 min and then resuspended and incubated overnight at room temperature in DMEM/F12 cell culture medium (ThermoFisher) containing 2 mM NaOH and 1% penicillin-streptomycin. The next day, microgels were homogenized once more at 15000 rpm for 3 min and centrifuged at 18500 G for 10 min. The supernatant was removed to leave a pellet of jam-packed alginate microgels that was stored at 4°C. One day before printing, the pellet was resuspended in double volume of fresh growth medium (described below) containing 4% HEPES (1M) and 4% NaHCO_3_ (37 g L^-1^ in sterile water, pH adjusted to 9.5 using NaOH) and incubated overnight. Microgel suspension was centrifuged at 18500 G for 10 min and supernatant removed. To generate SHAPE printing support, jam-packed microgels in the pellet were mixed with collagen solution (5 mg mL^-1^ Cultrex rat collagen I, R&D Systems) in 2:1 ratio. Collagen concentration was previously adjusted with growth medium to reach the final concentration of 1 mg mL^-1^ in the printing support. When cells were printed, SHAPE material was also supplemented with 2 μg mL^-1^ laminin (LN111, Biolamina), 5 μg mL^-1^ fibronectin (bovine, ThermoFisher), and 100 μg mL^-1^ hyaluronic acid (ab143634, Abcam).

### Rheology

Rheological characterization was performed on a Discovery HR-2 rheometer (TA Instruments) mounted with 25 mm parallel-plate geometry with the gap set to 0.9 mm. Fine-grain sandpaper was attached to the geometry in order to prevent slipping. Measurements were taken at 4°C on freshly prepared support material. In order to prevent loss of liquid due to evaporation in the temperature ramp experiment, the measuring cell was enclosed in a chamber with a liquid reservoir.

### Cell culture, 3D printing, and stem cell differentiation

Prior to printing, hNSCs (cell line named hVM1-Bcl-XL) were maintained as described before ^[32]^. In short, cells were maintained in flasks coated with Geltrex (ThermoFisher) in DMEM/F12 with GlutaMAX (ThermoFisher). Additionally, the medium contained 30 mM glucose, 5 μM HEPES, 0.5% w/v AlbuMAX (ThermoFisher Scientific), 40 μM each of L-alanine, L-asparagine monohydrate, L-aspartic acid, L-glutamin acid, and L-proline, 1% N2 supplement (ThermoFisher Scientific), 1% penicillin-streptomycin, 20 ng L^−1^ each of epidermal growth factor (EGF) and fibroblast growth factor (FGF) (R&D systems). For live imaging experiments, cells were labelled with 2 μM Calcein AM, 2 μM Calcein Red-Orange AM, or 10 μg mL^−1^ Hoechst 33342. Before printing, cells were enzymatically dissociated using trypsin and resuspended in growth medium with 0.1% xanthan gum (to prevent sedimentation during printing) at ~9×10^6^ cells mL^−1^ and loaded in a 500 μL Hamilton syringe. For the macroscopic prints in Fig. 2B, the ink was supplemented with 1 μm blue and red polystyrene microspheres (SigmaAldrich). Syringe was inserted in a volumetric extrusion printing head on a 3D Discovery bioprinter (RegenHU). Printing path was defined in BioCad software and cells extruded through a 27 G blunt needle inside SHAPE printing support kept at ~4°C (in a tissue culture well-plate or a custom container). Extrusion rate of 3.6 μL min^-1^ and lateral speed of 0.3 mm s^-1^ were used unless otherwise specified. After printing, the support was annealed at 37°C in a cell culture incubator for 30 min and then growth medium was added on top of annealed hydrogel. The next day, growth medium was replaced with differentiation medium that contains the same components as the growth medium with the exception of FGF and EGF that are replaced with 1 mM N^6^,2-O-dibutyryladenosine 3’,5’-cyclic monophosphate sodium salt and 2 ng L^-1^ GDNF. Differentiation medium was changed every second day.

### Immunocytochemistry

Prior to staining, printed constructs were fixed with 4% paraformaldehyde overnight at 4°C. Incubation with the blocking solution (5% serum with 0.25% Triton X-100), primary, and secondary antibodies was performed overnight at 4°C with gentle shaking on an orbital shaker. The following primary antibody were used in this study: β-tubulin III (mouse, Sigma Aldrich T8660, 1:1000), TH (rabbit, PelFreez Biologi-cals P40101, 1:1000), GFAP (rabbit, Dako Z0334, 1:1000; mouse, Biolegend SM121, 1:500). For fluorescent stainings, antibodies labelled with Alexa 546 (goat anti-mouse, Thermofisher Scientific A-11030, 1:500), Alexa 647 (goat anti-rabbit, ThermoFisher Scientific A-21245, 1:500), Cy5 (donkey anti-rabbit, Jackson Immunoresearch, 711605152, 1:500) were used. Nuclei were counterstained using DAPI (4’,6-diamidino-2-phenylindole) (1:1000).

### Microscopy

Fluorescence imaging of both live cells and immunolabelled printed constructs was performed on an inverted Ti2 microscope (Nikon) equipped with a CSU-W1 spinning disc system (Yokogawa) and a sCMOS camera (Teledyne Photometrics). 4x and 20x objectives were used. Z-stacks were acquired and tile-scans for larger samples. Reflectance microscopy was performed as follows. Samples were scanned at 647 nm excitation on a Nikon A1RHD confocal with a 100x Apo TIRF NA 1.49 objective, a non-polarizing 80:20 beam splitter was used to collect the reflected light from the sample. Images were processed in ImageJ (NIH).

### qRT-PCR

Prior to RNA isolation, printed constructs were incubated for 20 min in 5 mg mL^-1^ collagenase type I (STEMCELL Technologies) in PBS supplemented with 5% FBS and then lysed with RLT buffer. RNA isolation was performed using RNeasy Micro Kit (QIAGEN) according to manufacturer instructions. Approximately 1 μg of RNA was reverse transcribed using Maxima First Strand cDNA Synthesis Kit (ThermoFisher). cDNA was prepared together with SYBR Green Master mix (Roche) using Bravo liquid handling platform (Agilent) and analyzed by quantitative PCR on a LightCycler 480 II instrument (Roche) using a 2-step protocol. All qRT-PCR samples were run in technical triplicates, analyzed with the ΔΔCt-method, and normalized against the two housekeeping genes actin-beta (ACTB) and glyceraldehyde-3-phosphate dehydrogenase (GAPDH).

### Calcium imaging

Constructs were incubated for 1 hour at 37°C with 3 mM Fluo3 AM calcium indicator (ThermoFisher Scientific) in differentiation medium containing 0.02% Pluronic F-127 (Sigma Aldrich). Cells were then rinsed twice and incubated for 30 min at 37°C before imaging. Imaging was performed on the above-mentioned Ti2 microscope using a 20x objective. Exposure time was set to 50 ms. An environmental control chamber was used to maintain the temperature at 37°C and CO_2_ level at 5% during imaging. Images were analyzed in ImageJ (NIH) and plotted in Prism (GraphPad).

### Whole-Cell Patch-Clamp Recordings

Cellular constructs were transferred to recording chamber containing Krebs solution gassed with 95% O_2_ and 5% CO_2_ at room temperature and exchanged every 10-15 min during recordings. Krebs solution was composed of (in mM): 119 NaCl, 2.5 KCl, 1.3 MgSO_4_, 2.5 CaCl_2_, 25 glucose, and 26 NaHCO_3_. Micropipettes were generated using borosilicate glass capillaries with final resistance of 6 – 7 MΩ and filled with the following intracellular solution (in mM): 122.5 potassium gluconate, 12.5 KCl, 0.2 EGTA, 10 HEPES, 2 MgATP, 0.3 Na_3_GTP, and 8 NaCl adjusted to pH 7.3. For recordings, a Multiclamp 700B Microelectrode Amplifier (Molecular Devices) was used and data acquisition was performed with pCLAMP 10.2 software (Molecular Devices); current was filtered at 0.1 kHz and digitized at 2 kHz. Cells with neuronal morphology and round cell body positive for synapsin-GFP were selected for recordings. Resting membrane potentials were monitored immediately after membrane breaking-in. Thereafter, cells were kept at a membrane potential of −70 mV, and 500 ms currents were injected from −85 pA to +165 pA with 20 pA increments to induce action potentials. For inward sodium and delayed rectifying potassium current measurements, cells were clamped at −70 mV and voltage-depolarizing steps were delivered for 100 ms at 10 mV increments.

### Oxygen microsensor integration and 3D oxygen mapping

The technical aspects of the 3D oxygen sensor method, along with sensor integration into cell laden hydrogels, microscopy read-out, and signal processing have been described in detail previously ^[34]^. Here, we present a new application of the method to embedded to 3D extrusion printed tissue constructs. Briefly, oxygen sensor microbeads (CPOx Beads Red, Colibri Photonics GmbH) were embedded in SHAPE printing support either by direct extrusion printing of beads in along a preprogrammed path (before cells were printed) at a concentration of 2 mg mL^-1^ or by mixing beads with the cellular ink at a concentration of 0.7 mg mL^-1^. Direct printing of beads ensures placement of sensors in defined areas within and around tissue constructs. Beads were printed at extrusion rate of 0.2 μl s^-1^ and lateral speed of 1 mm s^-1^. Extrusion rates for the printing of cellular inks containing beads are indicated in the figures. For live-staining, cells were incubated for 3 h at 37°C in 6 μg mL^-1^ Calcein-AM (concentration in medium above granular hydrogel). Co-localized oxygen and live-cell maps were read-out with a confocal fluorescence and phosphorescence lifetime microscope (DCS 120 Flim System, Becker & Hickl GmbH) and image processing script was developed and executed in Matlab (Mathworks).

### Sacrificial ink printing and evacuation

Sacrificial ink was formulated by dissolving gelatin powder (bovine skin, Sigma Aldrich) in sterile water at 10% w/v concentration by constant mixing at 60°C under sterile conditions. Before printing, gelatin solution was liquified by heating and loaded into a 1 ml glass syringe (Hamilton). Printing was performed through a 21 G (500 μm inner diameter) blunt metal needle with extrusion rate of 22 μL min^-1^ and lateral speed of 0.6 mm s^-1^. Printing was performed in custom containers mounted on a glass slide with defined inlets and outlets. The containers were manufactured from Dental LT clear resin using stereolithography Form 2 printer (FormLabs) according to a CAD design created in Fusion 360 software (Autodesk). During printing, the syringe was heated to 50°C to liquify gelatin and allow smooth extrusion into SHAPE support. When deposited into the printing support, gelatin rapidly solidifies creating the sacrificial template for channel generation. After printing, the composite support containing the printed construct was incubated at 37°C overnight to anneal the support and liquify the sacrificial ink. Liquified gelatin structure was evacuated by injecting warm cell culture media. Food dye and colored polystyrene microspheres (SigmaAldrich) were added for better visualization of the perfused channels.

## General

Lund University Bioimaging Centre (LBIC) is gratefully acknowledged for providing experimental resources for reflectance microscopy.

## Author contributions

J.E, A.H. and J.K designed the project. J.K. carried out the main body of the experiments. M.F.W. and N.B.L. developed the oxygen sensing methodology. M.B. performed electrophysiology with the help of D.R.O. T.M. and J.L. provided materials and expertise for microparticle generation experiments. F.R.K. optimized microparticle generation methodology. A.M.S. supplied hNSCs and provided expertise on the experiments with hNSCs. M.P. contributed reagents and provided expertise on the evaluation of neuronal populations. T.E.A. and J.U.L. provided reagents and guidance for 3D printing experiments. J.K., M.F.W., and M.B. analyzed the experimental data. J.K. wrote the manuscript with the input from all authors.

## Data and materials availability

The data presented here are available upon request.

