## Supplementary materials for "Embedded 3D printing in self-healing annealable composites for precise patterning of functionally mature human neural constructs"

**This PDF file includes:**

Figs. S1 to S4
Movies S1 to S3

**Other Supplementary Materials for this manuscript include the following:**
Movies S1 to S3

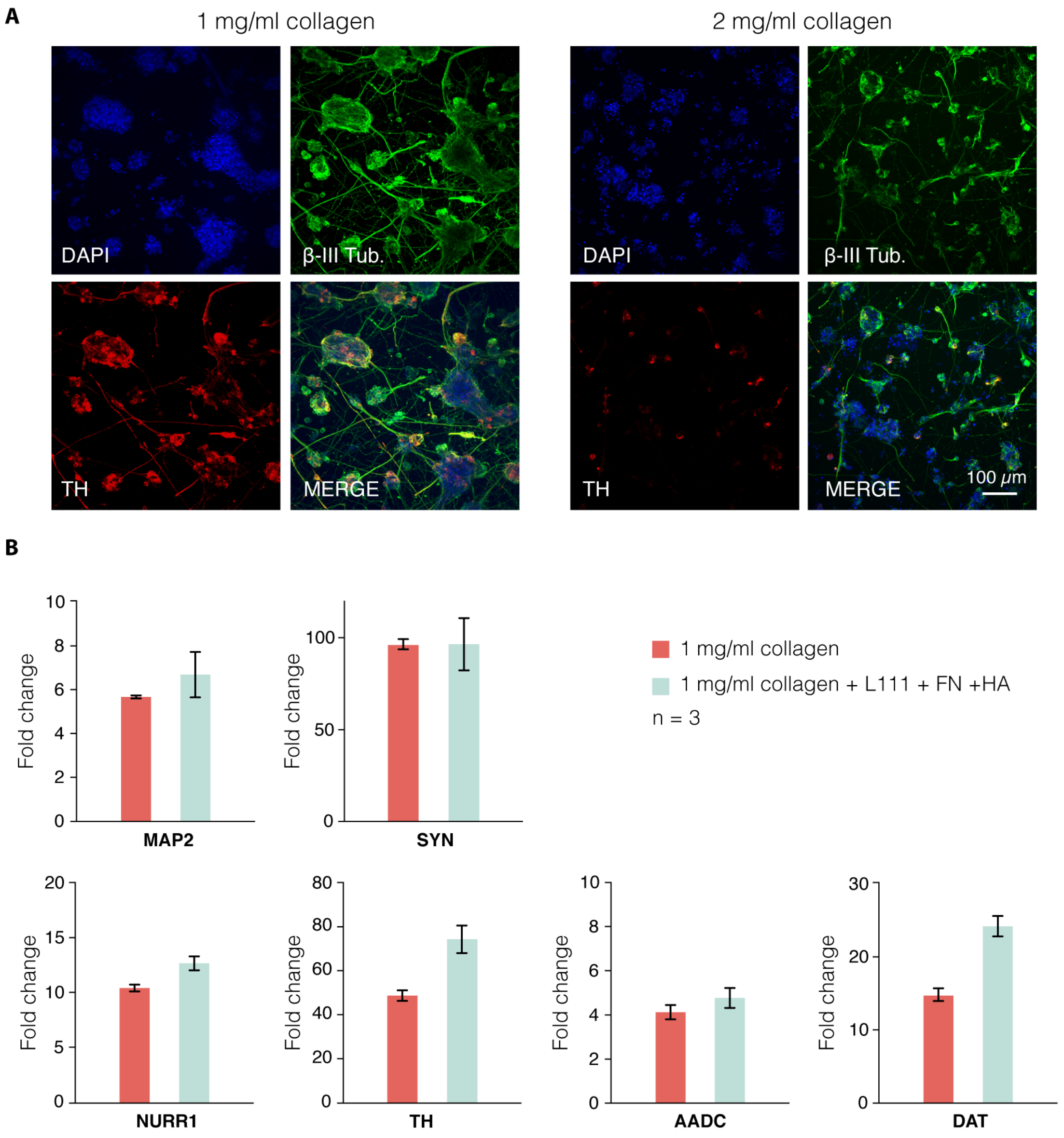


**Fig. S1 Soft ECM-mimicking hydrogels promote hNSC differentiation into ventral midbrain neurons.** (**A**) hNSCs were embedded in collagen hydrogels with varying collagen concentration (0.5 mg/ml, 1 mg/ml, 2 mg/ml). The lowest concentration did not crosslink sufficiently to provide 3D support to the cells. Immunocytochemistry after 4 weeks of differentiation indicates that softer hydrogels (1 mg/ml) provide a more permissive environment for expansion of cellular aggregates and growth and pathfinding of neuronal projections into the bulk of the hydrogel. Maximum intensity projections of fluorescence confocal images (150 μm thick optical slice) show that more extensive neuronal networks are formed in softer hydrogels marked by neuronal marker β-tubulin III, and higher expression of tyrosine hydroxylase (TH), a marker for dopaminergic neurons. (**B**) qRT-PCR gene expression analysis of hNSC differentiated for 4 weeks in collagen hydrogels ($2 \times 10^6$ cells/ml, 1 mg/ml collagen type I) with or without addition of other ECM components (2 μg/ml laminin 111, 5 μg/ml fibronectin, 100 μg/ml hyaluronic acid). While no significant difference between the two conditions was observed in the expression of neural markers, additional

ECM components cause significant increase in expression of genes related to midbrain identity (NURR1, AADC, TH, DAT). Non-differentiated hNSCs were used as a reference.

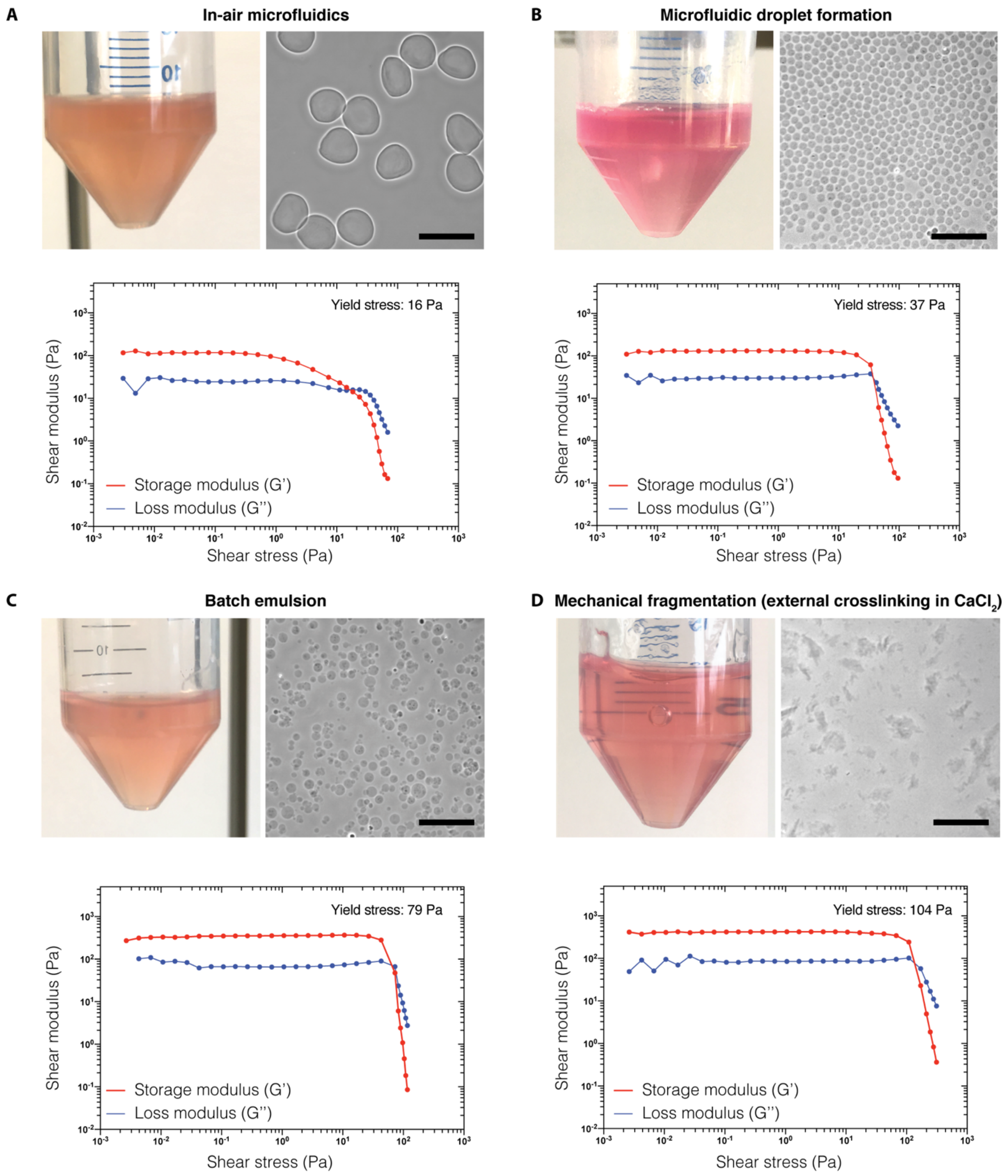


**Fig. S2. Alternative approaches for alginate microgel fabrication.** Four different ways of generating alginate microparticles is presented in each panel with a photograph of particle slurry in the jam-packed state (showing transparency of the material), phase contrast microscopy image (showing particle shape and size distribution), and measurement of shear moduli as a function of applied strain amplitude (showing that each jam-packed slurry undergoes shear stress-induced solid-to-liquid transition). (**A**) Microparticles generated using in-air microfluidics (supplied by IamFluidics, The Netherlands). (**B**) Microgels generated using flow focusing method in a PDMS microfluidic device (designed and fabricated in-house) where alginate solution (0.5 mg/ml, dispersed phase phase) with 1 mg/ml $CaCO_3$ was injected perpendicular to two streams of hexadecane (oil, continuous phase). The instabilities caused by surface tension led to droplet formation. Delayed addition of acetic acid (1:500 in hexadecane) to the flow after droplet formation ensures robust crosslinking of alginate hydrogels. The approach provides microgels with low polydispersity (~18 μm in diameter) but at low throughput. (**C**) Homogenization of alginate-in-oil emulsion allows higher throughput but without stringent control over microgel size. Alginate solution (0.5 mg/ml with 1 mg/ml $CaCo_3$) and hexadecane (1:3 ratio) were homogenized at 6000 rpm for 5 min. Then, same volume of. hexadecane with 0.4% acetic acid was added to the emulsion to initiate alginate crosslinking. (**D**) Mechanical fragmentation of externally crosslinked alginate hydrogels leads to flake-like microparticles with high polydispersity that, unlike the other three methods, results in highly transparent material when jam-packed. To generate these microparticles, 0.5% alginate solution was loaded into a syringe and dripped through a 500 μm ID needle into a 1% $CaCl_2$ bath thus creating a large amount of ~1 mm alginate beads that were then fragmented in a tabletop blender for 60 s. Scale bar 100 μm.

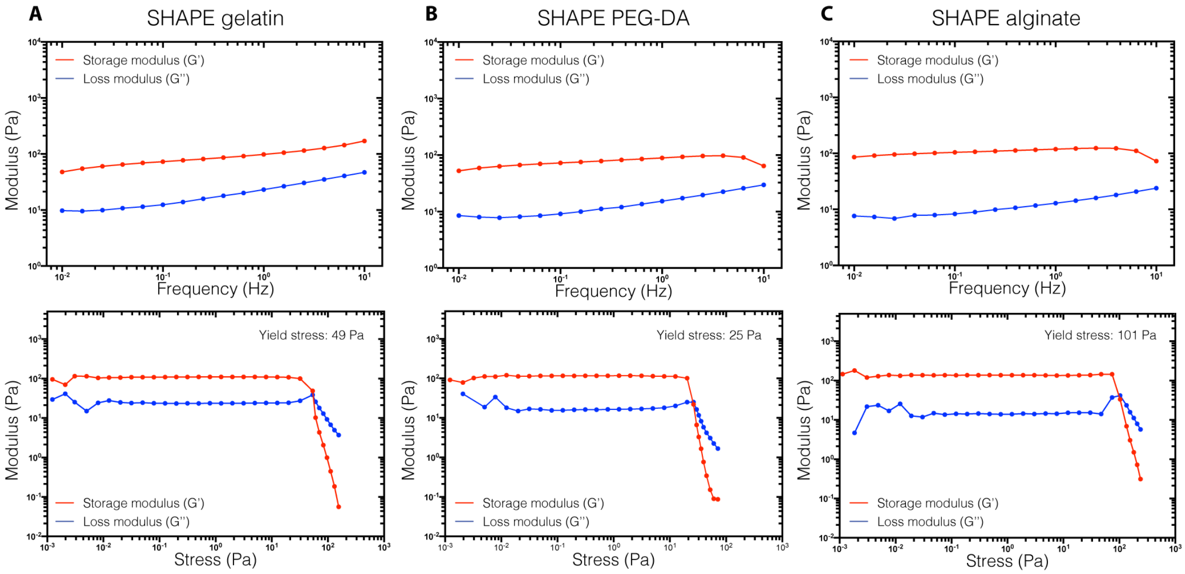


**Fig. S3. Rheological characterization of alternative SHAPE hydrogel formulations.** While keeping the same granular component (mechanically fragmented microgels from externally crosslinked alginate), we varied the contents of the continuous phase in order to showcase the versatility of the material landscape that can be explored for the generation of SHAPE hydrogels with different crosslinking mechanisms and for different biomanufacturing applications. In each column, the frequency sweep displayed in the top row shows that storage (G') and loss moduli (G") of alternative SHAPE hydrogel formulations are flat and show weak frequency dependence. Measurements of shear moduli as a function of applied strain amplitude (middle row) show solid-to-liquid transition. (**A**) Gelatin (10% w/v, bovine skin, Sigma Aldrich) was crosslinked enzymatically using microbial transglutaminase (Activa TI Transglutaminase, 10 U/ml in PBS) The crosslinking was performed overnight at 37 °C after which the crosslinked hydrogel was removed from the mold and enzyme washed away. (**B**) Poly(ethylene glycol) diacrylate (PEG-DA, 700 g/mol, 30% w/v, Sigma Aldrich) was crosslinked in the presence of photoinitiator (LAP, 0.5% w/v, Sigma Aldrich) using UV light with 365 nm wavelength in a curing chamber (CUREbox CB-4230, Wicked Engineering). After curing, the annealed construct was removed from the mold. (**C**) Alginate (1% w/v) was ionically crosslinked by adding 0.5% w/v $CaCl_2$ solution on top of the composite material and incubating overnight to allow $Ca^{2+}$ diffusion throughout the printing support. Crosslinking of alginate lead to ~50% shrinkage of the support.

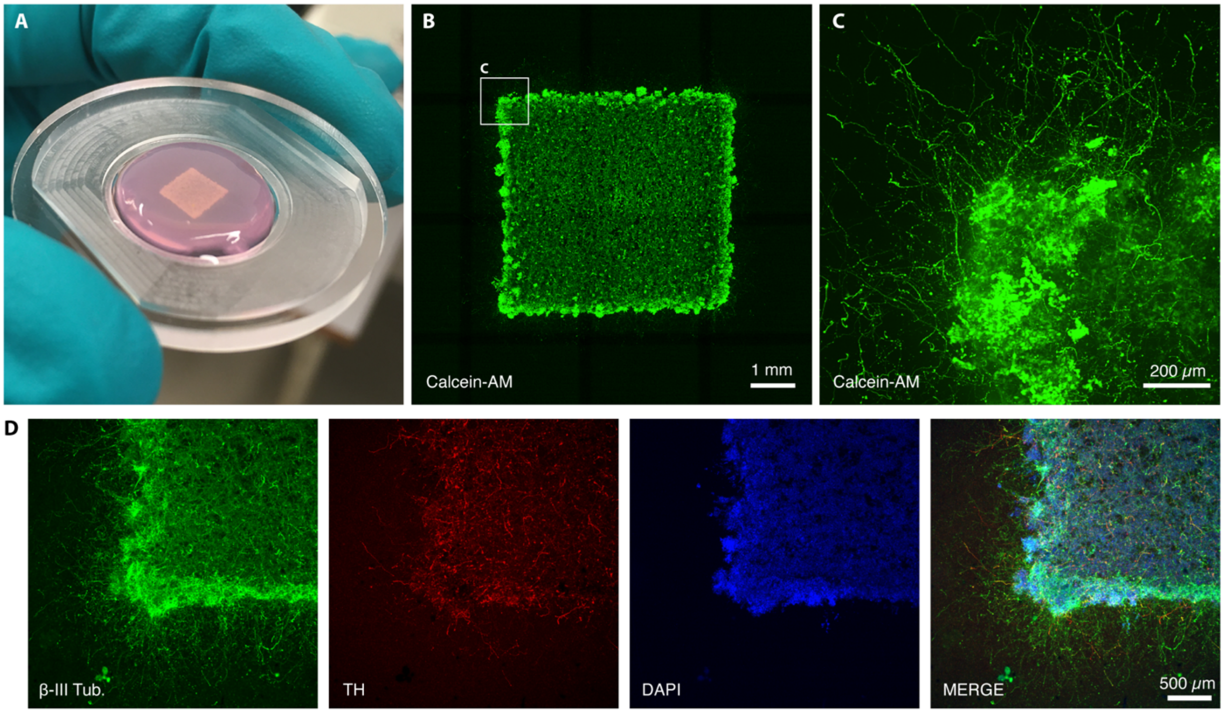


**Fig. S4. Generation of human dopaminergic neurons within 3D printed construct with dense infill.** (**A**) Photograph of a 3D printed square construct in annealed SHAPE hydrogel support after 2 months of differentiation. (**B**) Confocal image of the same construct labelled with live stain (Calcein-AM). (**C**) Maximum intensity projection (200 µm optical section) of a higher magnification fluorescence confocal image showing that neural projections extend away from the somatic volume and into the bulk of the support. (**D**) Fluorescence images of immunostained 3D printed construct after 2 months of differentiation displaying neuronal and dopaminergic markers (β-III tubulin and TH respectively).

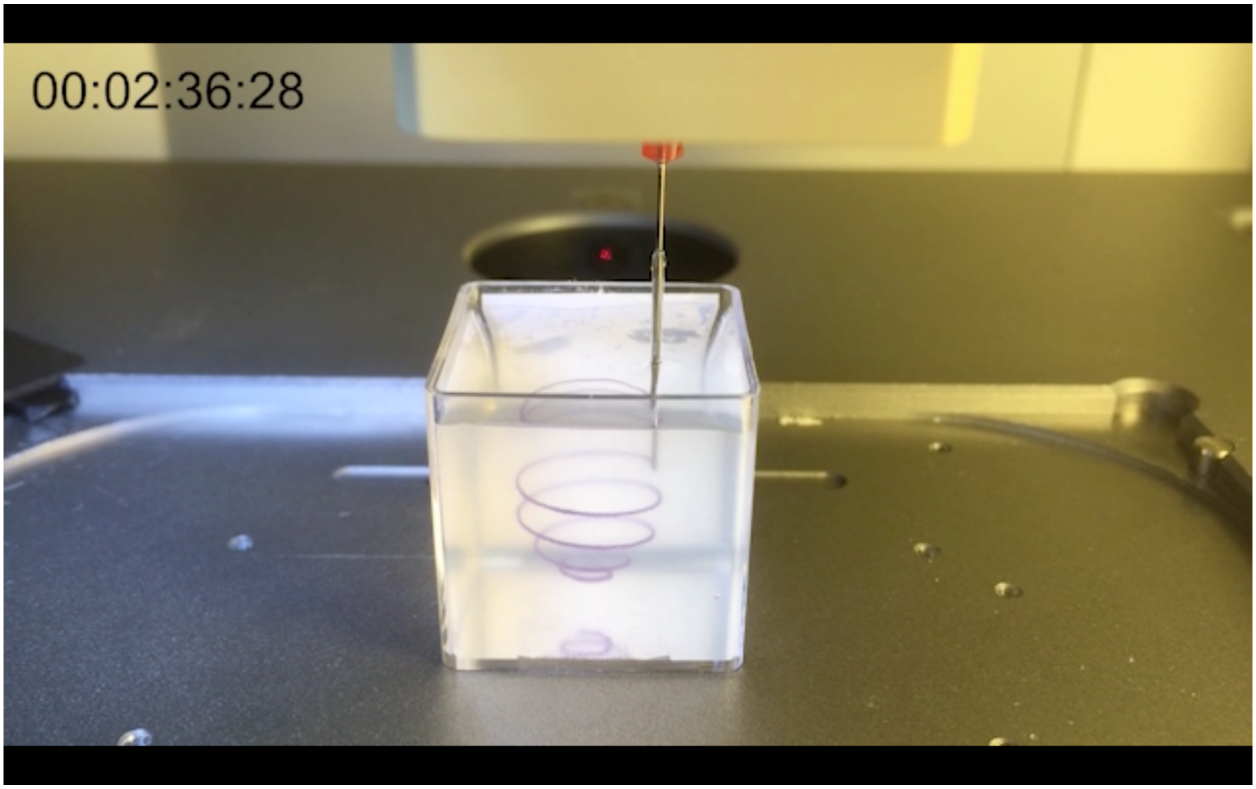


**Movie S1. Omnidirectional printing in SHAPE hydrogel support.** The movie shows the printing process of a spherical spiral using polystyrene beads as an ink to visualize the structure. Speed in the movie is increased to 11x. The width of the container is 25 mm.

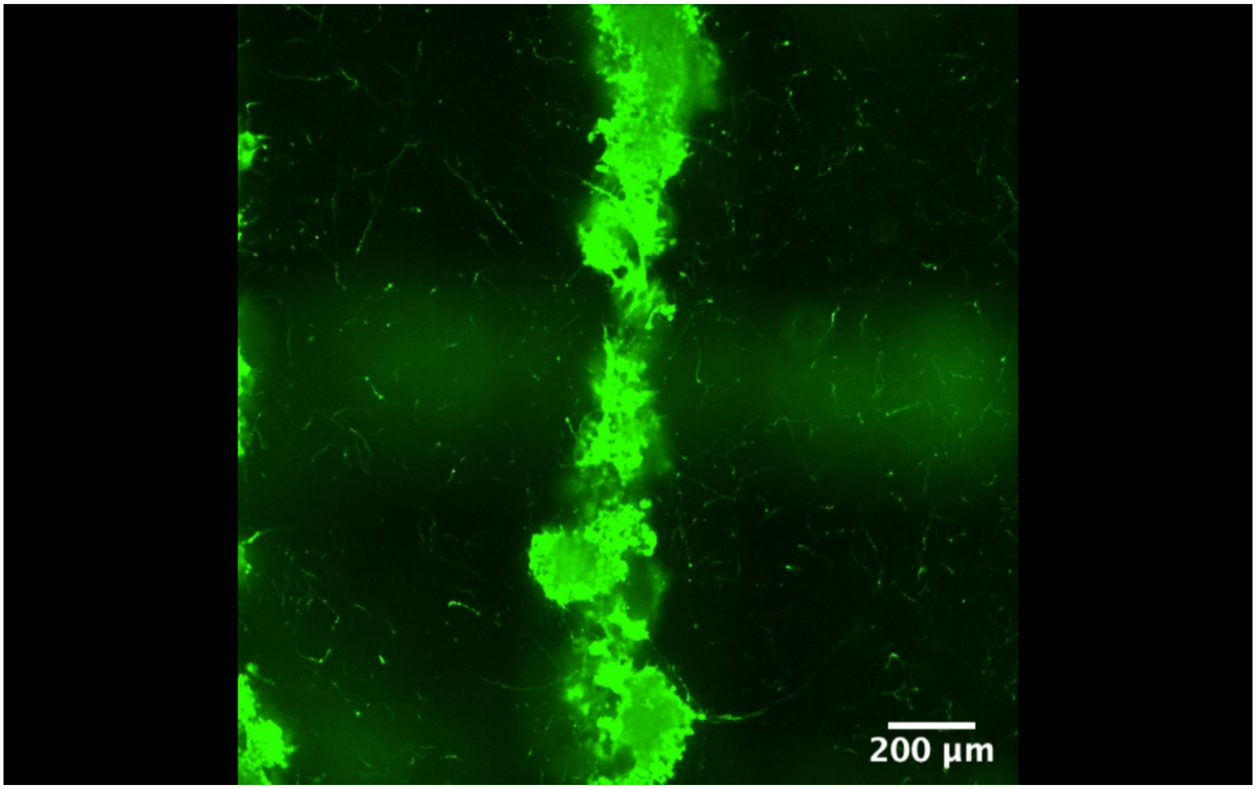


**Movie S2. Neuronal projections extend throughout the 3D space of SHAPE hydrogel.** The movie shows a walk-through along the z-axis for a stack of images (200 µm optical section) taken using a confocal microscope of neurons labelled with live stain (Calcein-AM) 2 moths after the structure was printed and hNSC differentiation started.

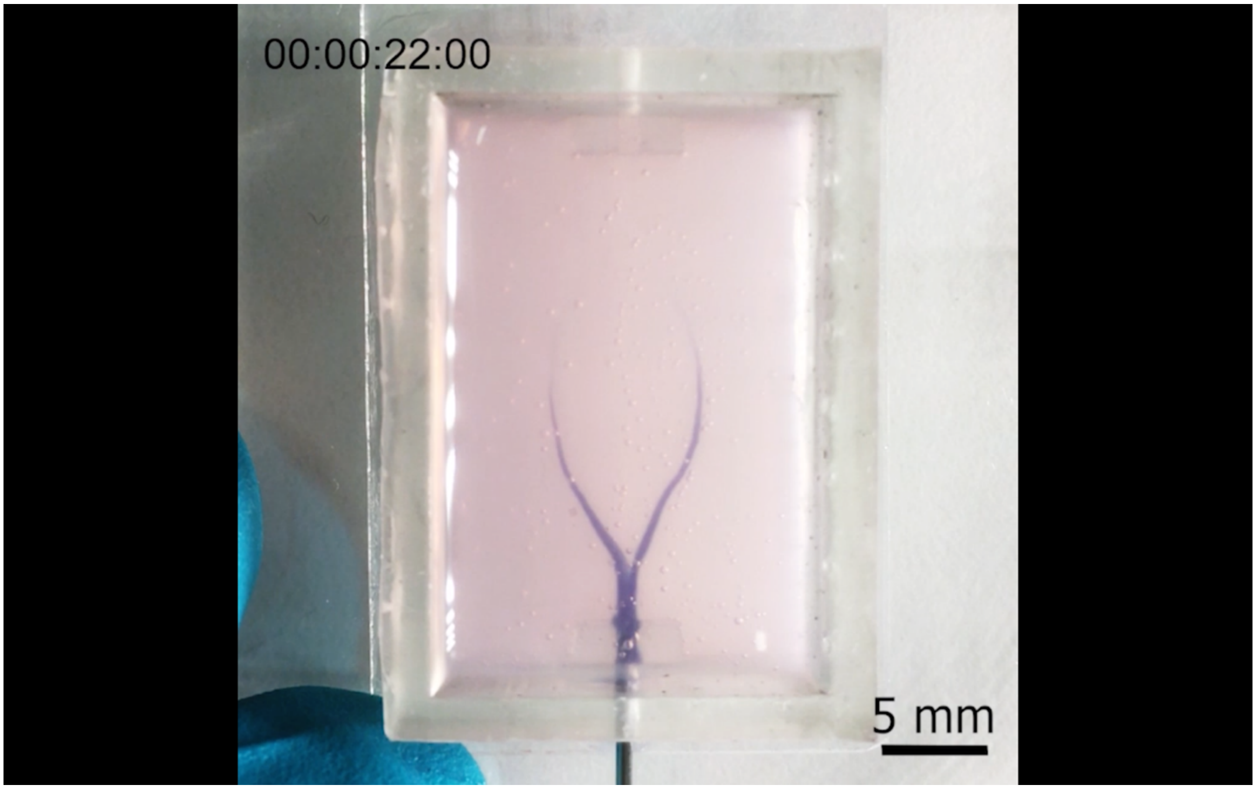


**Movie S3. Sacrificial ink displacement in a branching channel.** The movie shows displacement of a sacrificial gelatin ink from an annealed SHAPE hydrogel support. The material in channels is displaced simultaneously indicating successful strand fusion at the branching points during printing. Speed in the movie is increased to 6x.
